# Dehydration stress and Mayaro virus vector competence in *Aedes aegypti*

**DOI:** 10.1101/2023.05.08.539876

**Authors:** Jaime Manzano-Alvarez, Gerard Terradas, Christopher J. Holmes, Joshua B. Benoit, Jason L. Rasgon

## Abstract

The mosquito *Aedes aegypti* is a competent vector of multiple pathogens including dengue, Zika, yellow fever, chikungunya, and Mayaro viruses. *Ae. aegypti* is highly invasive and is currently present in the Americas, Oceania, Asia, and Europe, but its distribution and the pathogens it transmits are expected to change due to climate change. Relative humidity is an environmental variable that affects mosquito biology and distribution and can differ between location, habitat, and season, with mosquitoes facing significant variation in relative humidity during their lifespan. Low relative humidity can induce dehydration in mosquitoes, leading to alterations in physiological and behavioral responses relevant for pathogen transmission such as bloodfeeding and host-seeking behavior. In this study, we evaluated the short and long-term effects of dehydration stress on mortality and Mayaro virus vector competence in Ae. aegypti. Our results show that exposure to dehydration does not impact viral titers, nor infection, dissemination and transmission rates, in mosquitoes infected with Mayaro virus. However, we detected a significant effect of dehydration on mosquito mortality and blood feeding frequency regardless of infection status. The previously observed effects of higher feeding during dehydration and the current observation of altered survival along with no impact on vector competence suggest that the impact of dehydration on viral transmission in mosquitoes will likely be complex.

## Introduction

Vector-borne diseases (VBDs) are responsible for more than 17% of all reported infectious disease cases and cause ∼700,000 deaths worldwide (1). The mosquito Aedes aegypti, originally endemic to Africa, is now present worldwide and is a competent vector of many viruses including dengue, Zika, chikungunya, Mayaro, and yellow fever viruses (Reviewed in (2)). Dengue alone accounts for 2.3 million reported cases and over 1000 deaths in the Americas in 2013 (3). Although this mosquito species represents a significant public health threat, the association between climate and the pathogens that it transmits still requires further investigation (4).

Climatic factors such as precipitation, relative humidity, and temperature affect the distribution of mosquitoes and the pathogens they carry, and these variables are widely used for modeling VBD dynamics (5, 6). For example, decreases in relative humidity (RH) induce dehydration stress in the mosquito that alters its physiology and behavior, resulting in reductions in survival, nutrient reserves, oviposition and egg counts (7). Modeling studies suggest that environmental humidity is a driver of VBD occurrence due to the negative effect that dehydration in the mosquito has on vectorial capacity (6, 8). Nevertheless, we still require more empirical research to understand the effect of dehydration in vector-pathogen interactions.

RH has been found to be one of the determinants of *Ae. aegypti* distribution because the population of this species fluctuates depending on RH, along with precipitation and temperature (9–12). The current distribution of *Ae. aegypti* is already the widest ever recorded, and it is expected to further expand due to climate change (13, 14). RH is a fluctuating variable that can vary during the day; it has been reported under semi-field conditions that mosquitoes face RHs ranging from 50% to 100%, depending on the time of the day (15). Additionally, RH differs between indoor and outdoor settings (16, 17). Weather abnormalities can alter the environmental RH, such as is the case for dry heatwaves, which are periods of unusual hot weather characterized by an increase in temperature and decrease in RH (18). Heatwaves have economic and environmental impact worldwide, and their frequency is expected to increase due to climate change (19, 20). It is therefore expected that mosquitoes will face variable environmental RH during their lifespan, which is expected to have an impact of the survival of mosquitoes (21).

When RH decreases, mosquitoes invest energy in maintaining their osmotic balance to avoid dehydration through manipulation of their transpiration and evaporation rates or must respond to the physiological impact of water loss (6, 21, 23). Mosquitoes seek out beneficial microclimates, such as shrubs, to decrease water loss (24) and maintain osmotic balance by minimizing the water and ions loss through excretion using their highly efficient excretory system (21). When are continually exposed to arid conditions, they are able to change their cuticle composition and thickness (25, 26), and can induce morphological changes in their spiracles to avoid further water loss during severe dry seasons (27). In case they fail to maintain their osmotic balance over time, mosquitoes become more active and increase their host-seeking and blood feeding behaviors as an attempt to get needed water before dying of dehydration (6). If dehydration reaches critical levels, specific molecular changers occur to prevent and repair excessive damage, which includes the expression of antioxidants and heat shock proteins (6, 21-22).

Vector competence is the ability of a vector to become infected with a specific pathogen and transmit it to the next naive host during feeding (2). Environmental stressors, such as changes in temperature, have been previously shown to alter vector competence for alphaviruses and flaviviruses in *Ae. aegypti* (28–30). Because temperature and RH are closely related, we hypothesized that dehydration induced by different treatments of RH would affect viral vector competence of *Ae. aegypti* (vector) for Mayaro virus (MAYV; -L strain, pathogen). We show that exposure to RH stress affects mosquito mortality and bloodfeeding behavior, but we did not observe an effect on viral loads, nor infection, dissemination, or transmission rates (IR, DR, TR) of the virus in the mosquitoes.

## Methods

### Mosquito rearing

*Aedes aegypti* Liverpool strain mosquito eggs were originally provided by the NIH/NIAID Filariasis Research Reagent Resource Center for distribution by BEI Resources, NR-48921, NIAID, NIH. Insects were maintained and reared at the Penn State Millennium Sciences Complex insectary (University Park, USA) in 30×30×30cm cages, under 27°C±1°C, 12:12h light:dark cycle and 80% relative humidity. Larvae were fed koi pellets (Tetra Pond Koi Vibrance; Tetra, Melle, Germany). Adult mosquitoes were maintained with 10% sucrose solution *ad libitum*. For reproduction purposes, adult females were allowed to feed on anonymous human blood following a previously described membrane feeder protocol (31).

### Cells and virus stock

Vero cells (African green monkey kidney origin; CCL-81, ATCC, Manassas, VA, USA) were maintained in complete growth medium [Dulbecco’s modified Eagle’s medium (DMEM) supplemented with 10% fetal bovine serum (FBS) and 1% penicillin and streptomycin] in a 37 °C incubator with 5% CO_2_ [all reagents were purchased from Gibco, Thermo Fisher Scientific (Waltham, MA, USA)]. Mayaro virus genotype L strain BeAr505411 (BEI Resources, Manassas, VA, USA) was originally isolated from *Haemagogus janthinomys* mosquitoes in Para, Brazil in 1991. Virus was diluted in DMEM (multiplicity of infection of 0.01) and propagated in Vero cells for 1 hour, then cells were washed with DMEM and incubated with 30mL of complete growth medium for 24 hours. Then, infectious supernatants were aliquoted and stored at –80°C. Prior to their experimental use, viral titers of frozen stock aliquots were measured with focus forming assays (FFA).

### Humidity treatment setup

Three humidity treatments were prepared at a constant temperature of 27°C: 75%RH, 35%RH, and 80%RH (Control), which corresponds to regular insectary humidity conditions. To reach 75%RH and 35%RH humidity conditions, chambers were crafted with plastic transparent containers and cups holding supersaturated solutions of NaCl and MgCl_2_ respectively (32). Relative humidity was monitored before and during the experiment using 2 digital hygrometers (ThermoPro, Ontario, Canada) per chamber.

### Viral infection after exposure to dehydrating conditions

Three-to-five-day-old female mosquitoes were anesthetized with ice and sorted into three 20×30×20 board cages in groups of 120, then held for a day at normal insectary conditions to allow them to recover. The next day, mosquitoes were deprived of access to water, and cages were equally divided between the three humidity treatments for 18 hours of exposure (HE); number of dead mosquitoes per treatment was recorded at the end to calculate mortality rate. Mosquitoes were then fed for 1 hour on infected human blood spiked with infectious MAYV (1×10^7^ ffu/mL) at regular insectary conditions (27°C and 80%RH). An aliquot of infectious blood was collected, centrifuged, and stored at -80°C for further titration via FFA. Fully-engorged mosquitoes anesthetized on ice were sorted, placed in 10×10×10 cages and kept at 80%RH for the rest of the experiment. Blood-feeding rate, and daily survival were recorded. Vector competence assays were performed on twenty of the surviving mosquitoes per treatment at 7 and 14 dpi. Experiment was run in three biological replicates (Fig. 1a), and results are reported as the combination of them.

**Figure 1.**
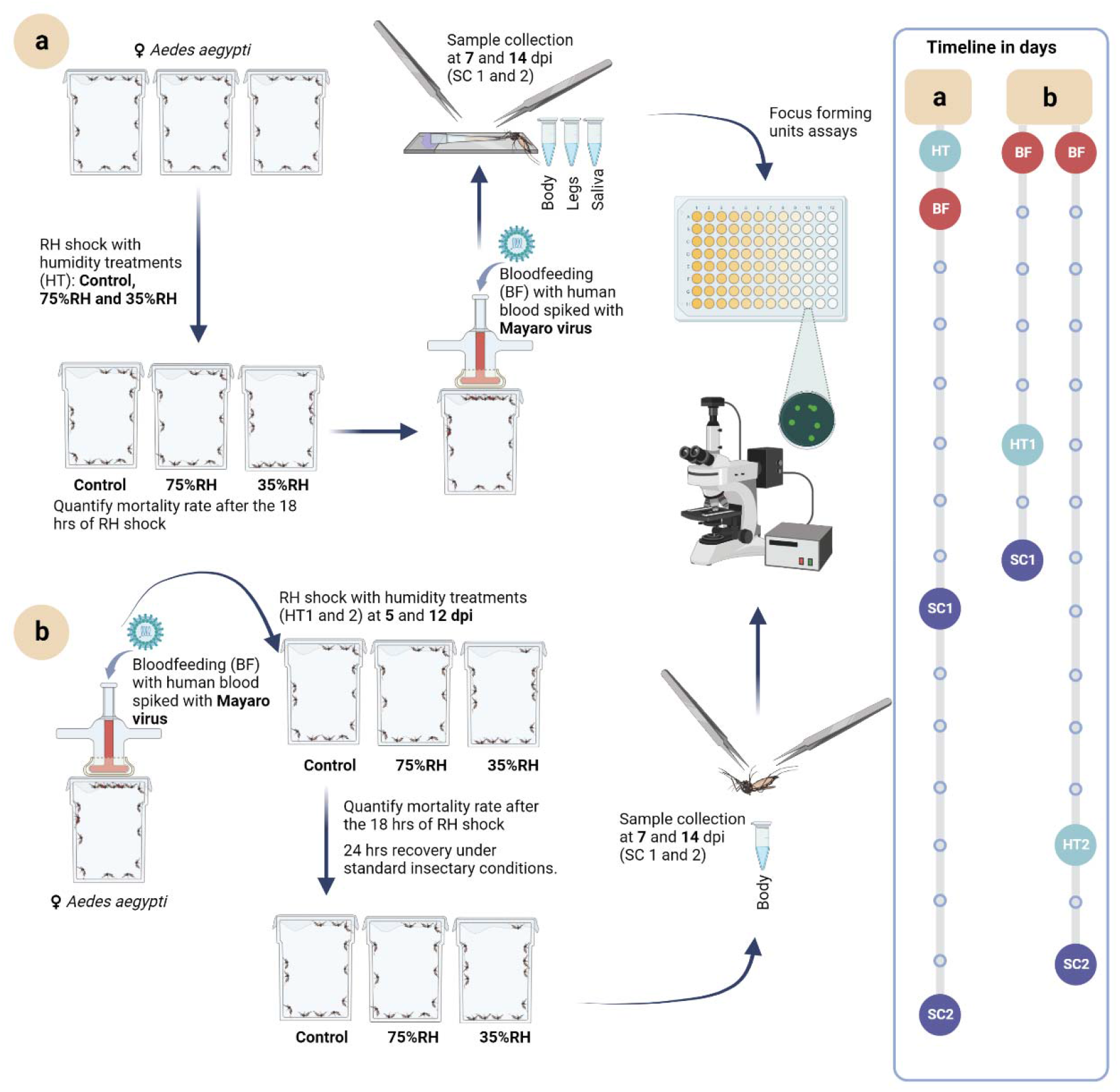
Pipeline for methods used in vector competence study. **a**. Long-term effect of 18 hours of exposure (HE) to dehydrating conditions (dehydration treatments). **b**. Short-term effect of 18 HE to dehydrating conditions. Mosquitoes did not have access to water nor sugar solution during varying RH exposure. The number of dead mosquitoes was counted just after the 18 HE. Once the exposure time was over, mosquitoes were put under standard insectary conditions (80%RH) with access to 10% sugar solution (Recovery conditions). Infectious bloodfeeding and focus forming assays were performed as in Brustolin et al., 2018. HT= Humidity treatments, BF= Bloodfeeding, SC= Sample collection.

### Viral infection before exposure to dehydrating conditions

Five to eight day-old female mosquitoes were anesthetized on ice and sorted into a 20×30×20 board cage containing 320 females, which were held for a day in regular insectary conditions (27°C and 80%RH). Mosquitoes were then fed for 1 hour on infected human blood spiked with infectious MAYV (1×10^7^ ffu/mL). An aliquot of infectious blood was collected, centrifuged, and stored at -80°C for further titration via FFA. Fully-engorged mosquitoes anesthetized on ice were sorted and placed in six 10×10×10 cm cages in groups of 40 and kept at regular insectary conditions. Three cages were randomly selected at 5 dpi and the other three at 12 dpi (for 7 and 14 dpi vector competence respectively). Selected boxes were then placed separately at the different humidity treatments for 18 HE. Mosquito mortality was assessed before and after the exposure to these humidity treatments. After 18 HE, mosquitoes were moved to recovery conditions for 24 hours to allow the virus replication. Next day, corresponding to 7 and 14 dpi, vector competence assays were performed in up to twenty of the surviving mosquitoes per treatment, collecting the whole body of mosquitoes in 300 μL of mosquito dilutant (20% heat-inactivated FBS in Dulbecco’s phosphate-buffered saline [PBS], 50μg/mL penicillin/streptomycin, 50 μg/mL gentamicin, and 2.5 μg/mL fungizone). Samples were quickly centrifuged and stored at -80°C until viral titering. Experiment was run in two biological replicates (Fig. 1b), and results are reported as the combination of them.

### Vector competence assays

At seven and fourteen days post-infection (dpi), mosquitoes were anesthetized with triethylamine (Sigma-Aldrich, St. Louis, MO, USA). A total of 20 mosquitoes were randomly selected per treatment and timepoint, when available. Legs were detached from the body and mosquitoes were forced to salivate into a pipette tip with a 1:1 mix of 50% sugar solution and FBS. Legs, body and saliva samples were collected in 2-mL safe-Lock tubes (Eppendorf, Hamburg, Germany) with 300, 300 and 100 μL of mosquito dilutant (see above) respectively, and placed on ice. Samples from body and legs were homogenized by a single zinc-plated, steel, 4.5-mm bead (Daisy, Rogers, AR, USA) using a TissueLyser II (QIAGEN GmbH, Hilden, Germany) on a 30Hz for 2 min cycle. Finally, samples were quickly centrifuged and stored at -80°C for further titration. Body, legs, and saliva samples were used to prepare 10-fold dilutions (10^2^ to 10^5^, 10^1^ to 10^4^, and 10^0^ to 10^1^ respectively) for the FFAs. Vector competence rates were reported as IR which stands for the proportion of infected bodies over the total, DR which stands for the proportion of infected legs over infected bodies, and TR which stands for the proportion of infected saliva over infected legs.

### Focus-forming assays (FFA)

Detection of infectious MAYV particles in samples from mosquito’s body, legs and saliva were carried out by FFAs in Vero cells. Vero cells were seeded in flat 96 well-plates at a density of 4×10^4 cells/well. The next day, series of 10-fold dilutions of the samples were prepared in FBS-free DMEM and 30 μL were used to infect the cells at 37°C and 5% CO_2_ for 1 hour. Then, supernatants were removed, replaced by 100 μL of overlay medium (1:1 mix of 1.6% methyl cellulose and complete growth medium), and incubated at 37°C and 5% CO_2_. The next day, cells were fixed with 4% paraformaldehyde for 15 min (Sigma, St. Louis, MO, USA), washed with PBS thoroughly, permeabilized with 0.2% Triton X in PBS for another 15 min, and washed with PBS. Viral antigens were detected using the primary monoclonal anti-chikungunya virus E2 envelope glyco-protein clone CHK-48 (BEI Resources, Manassas, VA, USA) diluted 1:500 in 1X PBS solution, as previously described (33). Primary antibody was recovered for further use, samples were washed with PBS, then primary antibody was marked with secondary antibody Alexa-488 goat anti-mouse IgG (Invitrogen, Eugene, OR, USA) at a 1:700 dilution with 1X PBS, followed by a last wash with PBS. An Olympus BX41 inverted microscope equipped with an UPlanFI 4X objective and a FITC filter was used for counting the MAYV foci.

### Statistical analysis and figure generation

Data were analyzed using R Studio (2023.3.0.386, PBC, Boston, MA, USA). Differences in infection rate (IR), (DR), (TR), and mortality and bloodfeeding rates were analyzed using Fisher’s test followed by multiple Bonferroni corrected comparisons. Since our data did not follow a normal distribution, Kruskal-Wallis test was used to compare viral titers in body, legs and saliva; the test was also used to assess differences among replicates. Survival curves were analyzed using a Log-rank test, which accounts for censored data. All p-values that were below 0.05 (p < 0.05) were considered significant. Graphs and plots were made with R Studio (2023.3.0.386, PBC, Boston, MA, USA), and Biorender.com. Final figures were assembled using Adobe Illustrator 2023 (27.4.1; Adobe, San Jose, CA, USA)

## Results

### Dehydration affects mortality and bloodfeeding in Ae. aegypti

Mosquitoes were exposed to RH treatments for 6, 12 and 18 hours of exposure (HE) to find the exposure levels that would allow mosquitoes to live in sufficient numbers to complete experiments. We detected 5% mortality under 35%RH conditions at 18 HE (Fig. S1), contrasting with the complete lack of mortality observed in the 75%RH and control (standard insectary RH; 80%RH) treatments. When mosquitoes were exposed for >18 hours, mortality significantly differed between treatments, dying at higher rates in the 35%RH (48%) compared to 2.5% in the 75%RH and 0% in the control treatment (Fig. S1). Since vector competence experiments required mosquitoes to survive for up to 14 days after being exposed to dehydration and virus infection, we chose 18 HE as dehydration time for the rest of the study.

To assess the interaction between relative humidity and mosquito biology, we equally distributed a total of 1065 mosquitoes into the three humidity treatments for 18HE, and measured mortality after exposure (Table 1). We found that after the RH treatments for 18 HE, mortality reached 4.5% in the 35%RH, significantly higher than 1.1% and 0.2% in 75%RH and control treatments, respectively (Table 1). Then, we challenged these same mosquitoes with MAYV-spiked blood, sorted bloodfed females and calculated bloodfeeding rates based on the proportion of mosquitoes that were observed to be engorged with blood over the total. We observed that the bloodfeeding rate was significantly higher in the 75%RH humidity treatment (Table 1).

**Table 1.**
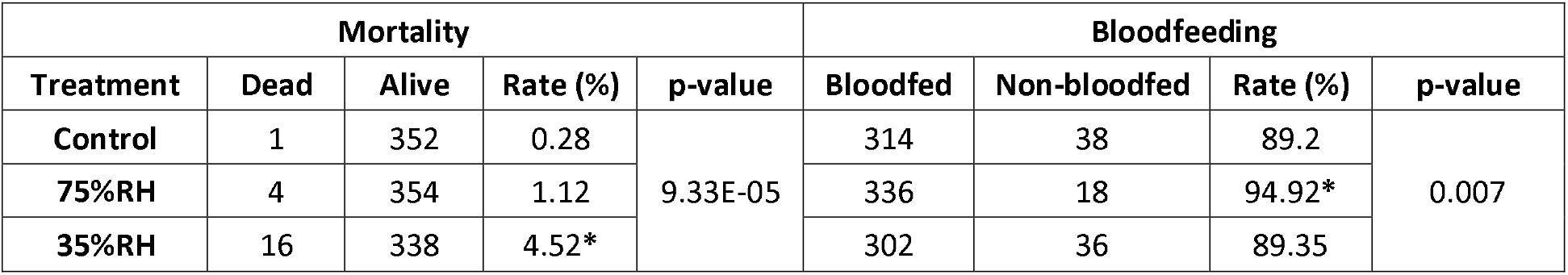
Mortality and bloodfeeding rates of naive mosquitoes exposed to 18 hours of RH that induce dehydration stress. * Indicates treatments that significantly differ from the other two. p-values were calculated with Fisher’s exact test followed by multiple comparisons analyses with Bonferroni correction.

### Dehydration stress does not affect long-term vector competence

To understand if the initial RH exposure alters the long-term survival and vector competence of mosquitoes challenged with virus, we returned mosquitoes to standard insectary conditions and vector competence was evaluated by dissecting relevant tissues at 7 and 14 dpi. When we compared viral titers of body, legs, and saliva, they were similar between RH treatments and replicates in all three tissues, showing no significant difference for either timepoint (Fig. 2a, b). Our results show that there is no difference in the IR, DR, and TR between treatments at both tested timepoints (Fig. 2c). Daily mosquito deaths were counted, and we used survival curves to assess differences between treatments. We found that there is no significant difference between the survival of mosquitoes for each dehydration treatment (Fig. 2d).

**Figure 2.**
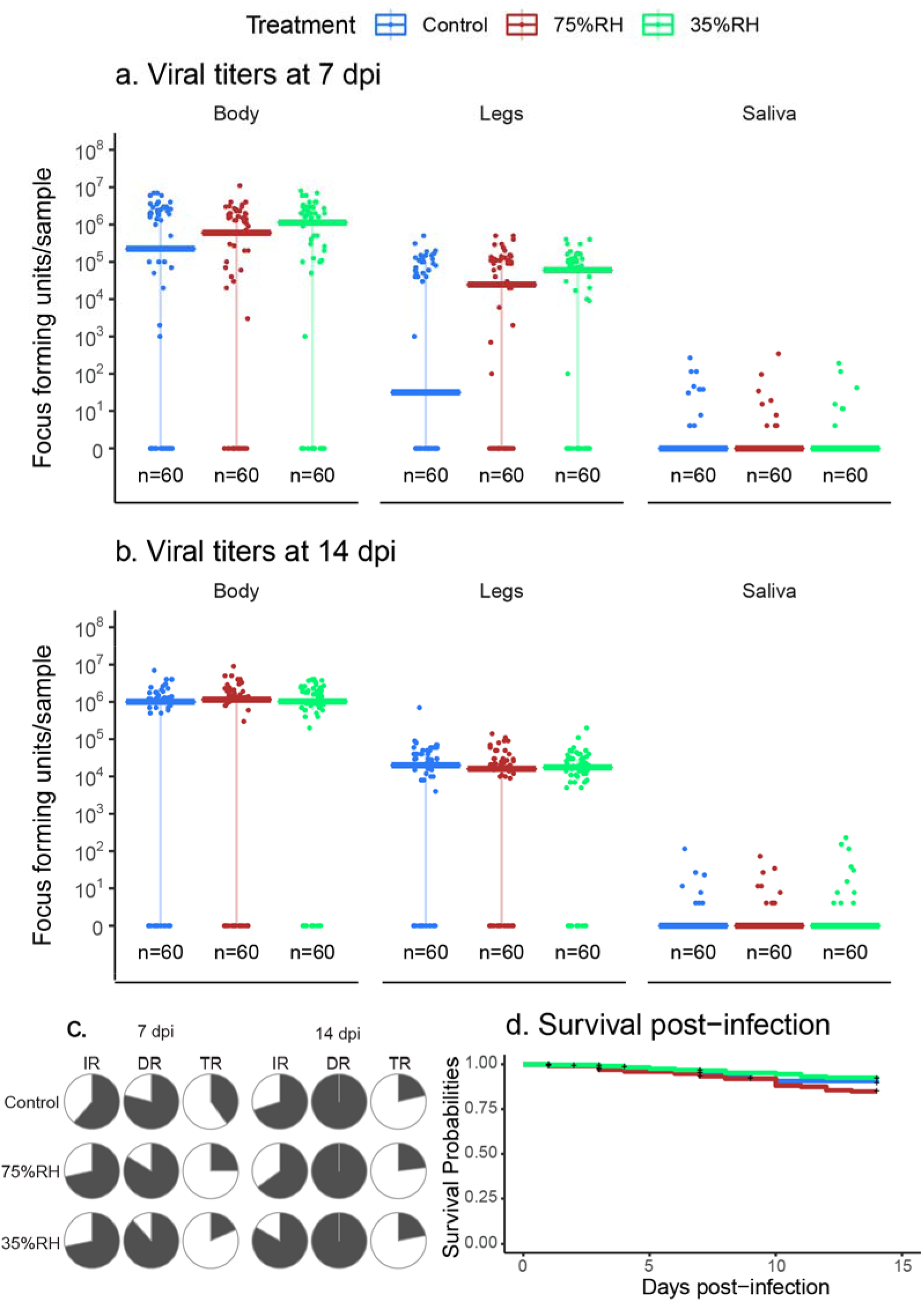
Vector competence at 7 and 14 dpi, and survival curve. Viral titers in mosquitos’ body, legs and saliva at 7 **(a)**, and 14 dpi **(b)**. n denotes sample size and bars represent the median, error bars represent data between the first and third quartiles. Virus concentration is presented on a logarithmic scale. **c**. Pie charts indicate infection (IR), dissemination (DR) and transmission rates (TR). **d**. Daily survival probabilities of *Aedes aegypti* after being exposed to dehydration and challenged with MAYV, + indicates when data was censored. We did not detect any statistically significant difference between treatments through the analyses (Kruskal-Wallis, Fisher exact test, and Log-rank test p-value > 0.05).

### RH shock prompts mortality in infected mosquitoes without affecting short-term vector competence

Since we did not observe altered vector competence due to long-term effects of dehydration, we aimed to understand the short-term effects in mosquitoes that were first infected with MAYV and then suffered dehydration (Fig. 1b). We challenged mosquitoes with MAYV, held them under standard insectary conditions, and then exposed them for 18 HE to the same three RH treatments at 5 and 12 dpi. These two timepoints were used because of viral competence collection timepoints at 7 and 14 dpi. Mortality rates were immediately recorded after the 18 HE (at 6 and 13 dpi). Our results (Table 2) show that mortality increases in the 35%RH treatment at 6 dpi, reaching a mortality of 42% that significantly differed from the 20% in the control treatment. Such difference between treatments is also found at 13 dpi, when the mortality in 35%RH treatment (47%) is significantly higher than the control (22%).

**Table 2.** Mortality rates of mosquitos exposed to 18 hours dehydration stress at 6 and 13 dpi. * indicates the treatment that significantly differs from the other two. *⍰indicates that the treatment significantly differs from the control. p-values were calculated with Fisher’s exact test, followed by multiple comparisons analyses with Bonferroni correction.

| Mortality |  |  |  |  |  |  |  |  |  |
| --- | --- | --- | --- | --- | --- | --- | --- | --- | --- |
| 6 dpi |  |  |  |  | 13 dpi |  |  |  |  |
| Treatment | Dead | Alive | Rate (%) | p-value | Treatment | Dead | Alive | Rate (%) | p-value |
| Control | 14 | 57 | 19.71 | 0.01 | Control | 14 | 50 | 21.88 | 4.42e-05 |
| 75%RH | 22 | 50 | 30.55 |  | 75%RH | 7 | 52 | 11.86 |  |
| 35%RH | 30 | 41 | 42.25*□ | | 35%RH | 30 | 34 | 46.88* | |

Once mortality was assessed, mosquitoes were allowed to recover for 24 hours under standard insectary conditions so the virus could resume replication. Instead of tissue dissections, we collected whole body samples immediately after the recovery period (at 7 and 14dpi) aiming to understand the short-term effect of dehydration on MAYV. Results are reported as viral titers in the bodies (Fig 3a) or prevalence of infection (IR, Fig 3b). Our results indicate that exposure to varying RH after viral challenge did not affect the IR nor the viral titers at either timepoint.

**Figure 3.**
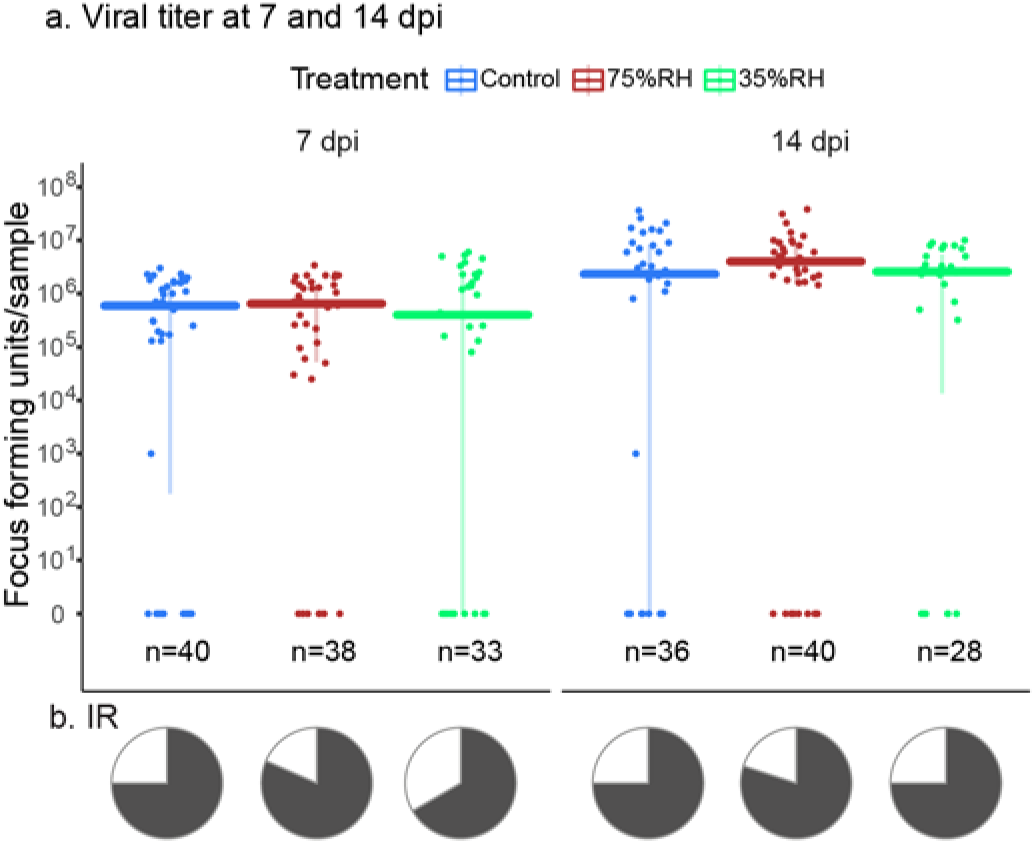
Vector competence of mosquitos exposed to humidity treatments after being challenged with MAYV. **a**. Data are shown as viral titers in mosquitos’ body at 7dpi and 14 dpi. n denotes sample size, bars represent the median, and error bars represent data between the first and third quartiles. Data is shown in logarithmic scale. **b**. Pie charts indicate infection rate (IR). We did not detect a statistically significant difference through the respective statistical analyses (Kruskal-Wallis and Fisher exact test p-value > 0.05)

## Discussion

Dehydration in mosquitoes occurs due to a combination of lack of access to water, increases in temperature, and decreases in environmental humidity (6). Mosquitoes can use multiple strategies to counter dehydration, such as resting in microhabitats with higher moisture, altering their activity patterns, and increasing bloodfeeding activity (reviewed in (7)). Under our experimental design, mosquitoes were exposed to dehydration without the possibility of using such strategies to curtail dehydration during exposure to low RH. We found that dehydrating *Ae. aegypti* mosquitoes at 75%RH is enough to increase bloodfeeding rates without compromising mortality, while exposure to low RH increased mortality, consistent with previous observations in *Culex pipiens* (6). These mortality effects were increased when mosquitoes were previously infected with MAYV, suggesting that viral infection may increase the sensitivity to dehydration stress in mosquitoes. This has been previously shown with other arboviruses, and stressors originated from environmental variables such as thermal stress (34– 36), likely due to the demand of resources for keeping cellular homeostasis, the cost of immune responses against the viral infection, and virus-mediated changes in gene expression (37–39) Importantly, dehydration can be extremely stressful and require specific factors to maintain cellular homeostasis and allow recovery that could very well be impaired during an active viral infection (6, 21-22). Additionally, we found that once mosquitoes survived dehydration and were placed under standard insectary conditions, they showed no difference in daily survival between treatments for a period of 14 days, supporting the theory that bloodfeeding allows mosquitoes to recover from dehydration (40) even when facing a newly acquired viral infection.

In this study, we tested the short- and long-term effects of dehydration stress over viral vector competence. In our case, we did not observe a change in IR, DR and TR when mosquitoes faced different levels of dehydration for a period of 18 HE (Fig 2abc, Fig. 3). There is a large body of literature showing that viral vector competence can be impaired under circumstances that stress the mosquito, including changes in environmental variables such as temperature (26–28, 39). Since dehydration induces physiological changes in the mosquito (21-22, 41-43), we hypothesized that dehydration would affect vector competence as well. However, it is possible that periods longer than 18 HE to low humidity conditions are required for dehydrating mosquitoes enough to affect their viral vector competence, which could be challenging considering the increase in mortality reported here. Repeated bouts of dehydration have been shown to directly impact mosquito physiology and reproduction (41), suggesting that future studies may want to target how chronic and repeated bouts of exposure to RH stress may impact viral transmission. Although dehydration did not affect vector competence in these experiments, it does not necessarily mean that dehydration does not affect transmission dynamics. Since mortality and density of the mosquitoes (vectors) affect arbovirus transmission (44, 45), it could be considered that a period of dehydration will influence the transmission dynamics in arboviruses by impacting other biological factors such as mortality and feeding rates, as previously modeled for *Culex pipiens* and West Nile virus (6).

MAYV is primarily transmitted by arboreal *Haemagogus* mosquitoes from non-human primates to humans in a sylvatic cycle (46), but maintenance of this mosquito species is challenging under laboratory conditions (47) and hinders vector-pathogen interaction studies. There is evidence of natural infection of MAYV in adult *Ae. aegypti* from Brazil (48), and is a competent vector of MAYV under laboratory conditions. Thus, *Ae. aegypti* stands as an important model species for the study of vector-MAYV interactions under laboratory conditions.

Several aspects that were not covered in this research should be considered for future studies. First, *Ae. aegypti* is present worldwide (14), and environmental variables such as RH, precipitation and temperature differ between locations and habitats. Mosquitoes distributed in dryer areas have been found to change their phenotype to decrease water loss (25, 26), so studying *Ae. aegypti* strains derived from different mesoclimates could be informative. As vectorial capacity relies heavily in the extrinsic incubation period (EIP) of the virus inside the mosquito (45), it will be relevant to assess how dehydration and RH might affect the length of EIP in further studies of vector competence. Recent comparative studies between *Ae. aegypti* populations have shown differences in feeding patterns and viral transmission in relation to environmental factors and urbanization (49-50), which suggest that dehydration and viral transmission dynamics may vary between *Ae. aegypti* lineages. MAYV has been shown to infect some species of *Anopheles* mosquitoes *in vivo* and *in vitro* (31, 51, 52). Several *Anopheles* species (that are also vectors of malaria) are distributed in the Americas (Reviewed in (53)), including countries where MAYV has been reported, such as Colombia, Venezuela and Brazil (54). Although some progress has been made to decipher how RH affects the biology of these mosquitoes (55-56), more research is still required. Thus, it would be important to explore how RH affects the mortality and vector-MAYV interactions in *Anopheles*.

In conclusion, our work suggests that dehydration can increase bloodfeeding behavior and mortality in mosquitoes, depending on the severity of dehydration. However, mosquitoes are able to recover from this state once RH increases to more stable levels and food sources become available. Additionally, under this experimental design, we found that dehydration did not play a role in driving the viral vector competence in Ae. aegypti. Finally, we suggest that further studies should explore the relationship between RH and vector competence (including EIP) in several viral strains and species of mosquitoes that are competent vectors of MAYV.

## Data availability

All data are available as supplementary material

## Acknowledgements

We thank Amelia Romo, Heather Engler and Kaylee Montanari for support in mosquito rearing, and Kristine Werling, Renuka Joseph, Hargobinder Kaur, Rachel S. Krizek and Sultan Asad for technical support.

This work was funded by NIH/NIAID grant R01AI148551 to JBB and JLR, and NIH/NIAID grants R01AI116636 and R01AI150251, USDA Hatch funds (Project #4769), a SEED grant from the Penn State Huck Institutes of the Life Sciences, and funds from the Dorothy Foehr Huck and J. Lloyd Huck endowment to JLR. JMA was supported by the Fulbright Pasaporte a la Ciencia program, a Colombia Científica component from ICETEX, in collaboration with Fulbright Colombia.

## Supplemental material

## Methods

### Assessment of mortality at different timepoints of exposure to relative humidity treatments

Three humidity treatments were prepared: 75% RH, 32%RH, and control treatment, the latter was set at regular insectary humidity conditions (80%RH). To reach 75% and 35% relative humidity conditions, chambers were crafted with plastic transparent containers, holding cups filled with supersaturated solutions of NaCL and MgCl_2_ in the inside respectively. Three- to five-days old female mosquitoes were anesthetized with ice and sorted into nine 20×30×20 board cages in groups of 120 individuals, then held for a day at normal insectary conditions to allow them to recover. The next day, mosquitoes were deprived of access to water, and cages were equally divided between the three humidity treatments. Mortality was recorded for all the cages every 6 hours, and one cage per treatment was selected for immediately feeding on human blood for 1 hour to compare blood-feeding rates. The cages that were selected in each timepoint were then excluded from the rest of the experiment. Two replicates were performed, the timepoints for each replicate were 6, 12 and 18; and 12, 18 and 24 hours of exposure (HE) respectively.

### Results

**Figure S1.**
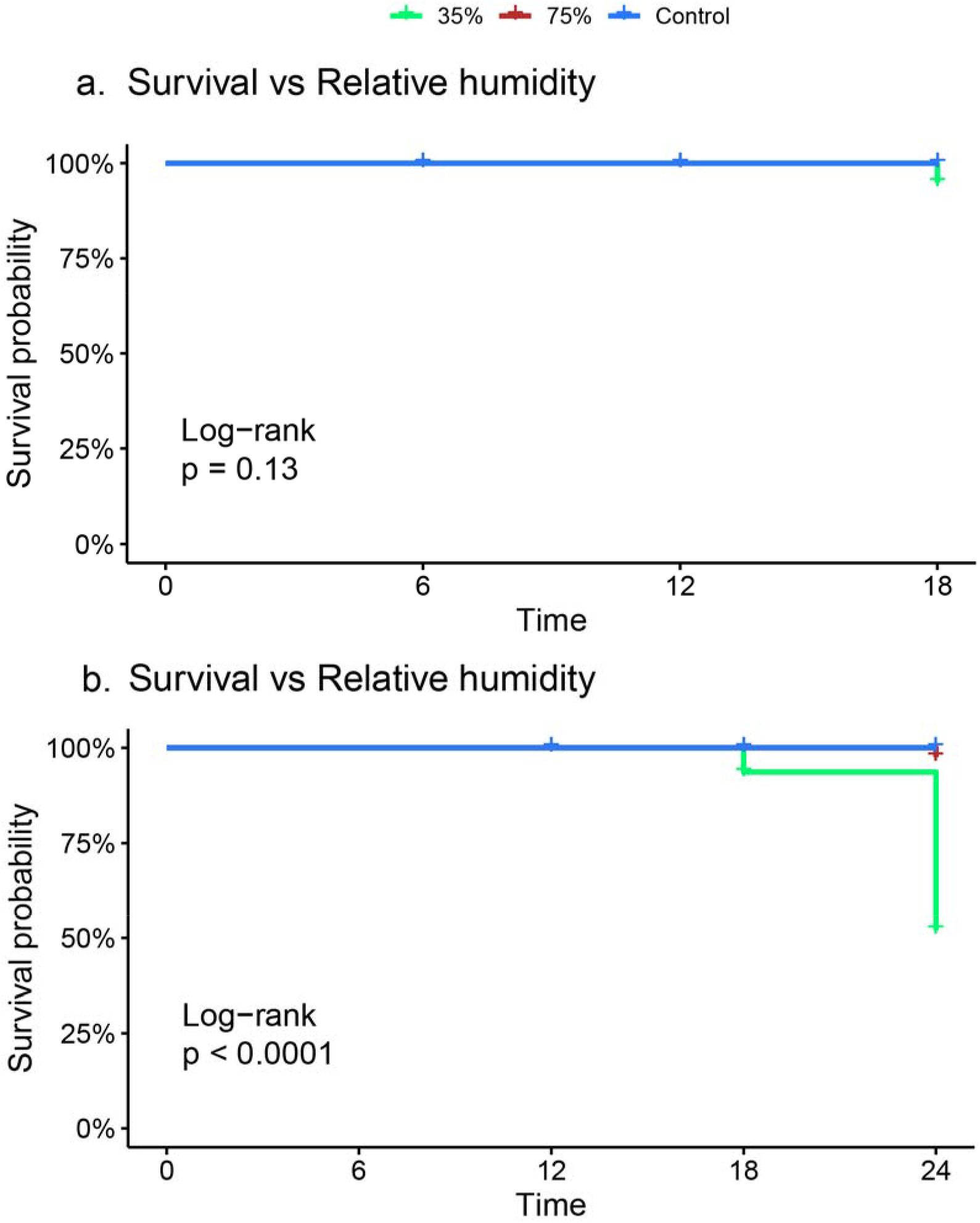
Mosquito mortality per HE to the humidity treatments. Each graph represents the survival curve of mosquitoes challenged with the temperature treatments for different times of exposure in 2 replicates. **a**. The first replicate tested the timepoints 6, 12 and 18 HE. **b**. Second replicate tested 12, 18 and 24 HE timepoints.

**Table S1.**
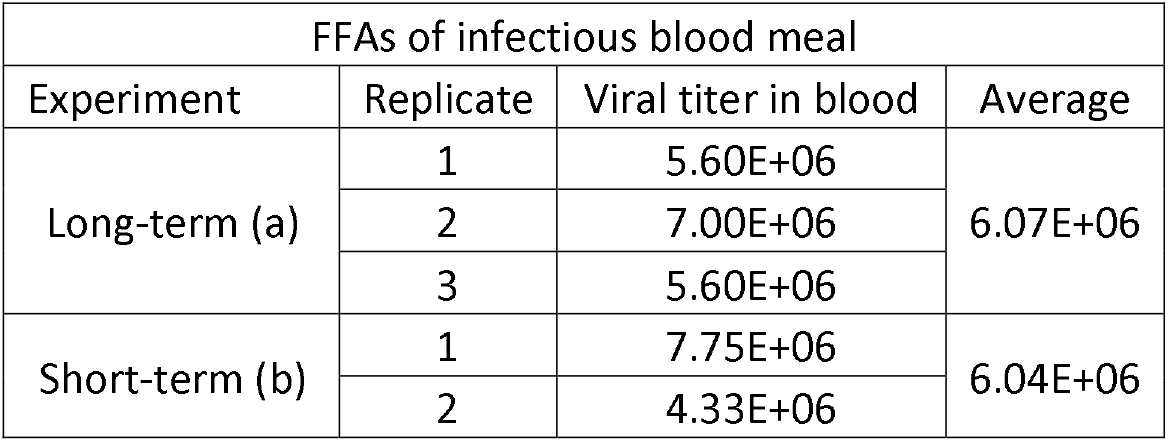
Viral tiers of aliquots of infectious blood offered to mosquitoes. Samples were stored in cold at -80°C, until virus tittering with FFA. Results differ from initial concentration (1E+07) because freezing and thawing the samples can decrease their viral loads.

